# Discovery of a Novel Non-MET-Mediated Otoprotective Compound Against Aminoglycoside-Induced Ototoxicity

**DOI:** 10.64898/2026.05.14.725303

**Authors:** Elizabeth Kara, Christopher Nicolet, Shaikh Emdadur Rahman, Travis Hudok, Caleb Leach, Kaden Falkner, Kenneth A. Cornell, Dong Xu

## Abstract

Aminoglycoside (AG) antibiotics are indispensable for treating severe infections but frequently cause irreversible hearing loss, with no approved preventive therapies. Using *in vivo* zebrafish lateral line screening combined with computational scaffold-hopping, we identified a novel class of otoprotective compounds. Starting from the ion channel modulator MR16728, we discovered compound 28510 as a potent lead compound. Compound 28510 provided robust, dose-dependent protection against AG-induced hair cell damage, restoring neuromast hair cell integrity to near-control levels in acute assays and demonstrating broad efficacy across clinically relevant AGs (gentamicin, tobramycin, amikacin, streptomycin) in chronic exposures. Importantly, 28510 exhibited a favorable therapeutic window, with low micromolar 50% hair cell protection concentration (HC_50_) values consistently below toxicity thresholds. Mechanistically, FM1-43 and Texas Red-conjugated gentamicin uptake assays revealed that 28510 does not inhibit mechanotransduction (MET) channel-mediated AG entry, distinguishing it from current clinical candidates and pointing to a novel intracellular protective mechanism. 28510 preserved AG antibacterial activity in *E. coli* assays, supporting its translational compatibility as a co-therapeutic agent. Combinations of 28510 with related analogs did not yield synergistic protection; 28510 alone remained the most effective compound. *In silico* absorption, distribution, metabolism, and excretion (ADME) predictions further confirmed its highly favorable drug-like properties, including excellent intestinal and oral absorption. Together, these findings establish 28510 as a first-in-class, non-MET-mediated otoprotective lead with broad efficacy and a favorable therapeutic profile, highlighting a new strategy for preventing AG-induced hearing loss.

## INTRODUCTION

Aminoglycoside (AG) antibiotics remain a cornerstone in the treatment of severe and life-threatening infections, particularly those caused by aerobic Gram-negative bacteria. Since the discovery of streptomycin in 1944, multiple AGs, including gentamicin, tobramycin, and amikacin, have been widely used in clinical settings for conditions such as sepsis, complicated urinary tract infections, cystic fibrosis–associated infections, and multidrug-resistant tuberculosis^1,2^. Despite the development of alternative antimicrobial classes, AGs continue to play an essential role due to their rapid bactericidal activity and synergistic effects when combined with other antibiotics^3^.

However, the clinical utility of AGs is significantly limited by their ototoxic side effects, which can result in permanent hearing loss and vestibular dysfunction. The incidence of AG-induced ototoxicity is estimated to affect approximately 20% of treated patients, with higher susceptibility observed in individuals harboring specific mitochondrial DNA mutations, such as the A1555G mutation in the 12S rRNA gene^4–6^. Mechanistically, AGs enter the inner ear through the blood–labyrinth barrier and accumulate in the endolymph, where they preferentially target sensory hair cells, particularly outer hair cells in the cochlea^7^. Entry into hair cells is predominantly mediated by mechanotransduction (MET) channels, although alternative pathways, including endocytosis and transient receptor potential (TRP) channels, have also been implicated^8^.

Once internalized, AGs disrupt intracellular homeostasis, triggering pathways that lead to hair cell death. These mechanisms include dysregulation of intracellular calcium signaling, mitochondrial calcium overload, and the generation of reactive oxygen species (ROS), ultimately activating apoptotic pathways^9–11^. Despite these established modes of ototoxicity, the precise mechanisms for each AG remain incompletely understood, limiting the development of effective strategies to prevent AG-induced hearing loss.

To date, most otoprotective strategies have focused on blocking AG entry into hair cells via MET channels. For example, ORC-13661, a small-molecule MET channel blocker, has demonstrated protective efficacy in preclinical models and is currently undergoing clinical evaluation^12,13^. However, such approaches raise concerns regarding potential interference with normal MET channels and auditory function, as well as incomplete protection due to alternative routes of AG entry.

Phenotypic drug discovery in *in vivo* models provides a powerful approach for identifying compounds with protective effects independent of predefined molecular targets. The zebrafish (*Danio rerio*) lateral line system is a well-established model for studying sensory hair cell biology and drug-induced ototoxicity, owing to its accessibility, structural and functional similarity to mammalian hair cells, and suitability for high-throughput screening^14–16^. Using this model, previous studies have identified ion channel modulators, such as MR16728, that confer partial protection against AG-induced hair cell damage; however, their therapeutic potential has been limited by cytotoxicity, hindering their advancement as viable drug candidates^17^.

In this study, we employed a combined *in vivo* phenotypic screening and computational scaffold-hopping strategy to identify novel otoprotective compounds with improved efficacy and safety profiles. Building on the structure of MR16728, we identified compound 28510 as a lead candidate with robust protective activity across multiple AGs and acute/chronic exposure assays. Notably, this compound does not block MET channel-mediated AG uptake, suggesting a previously unrecognized intracellular mechanism of protection. These findings highlight a new therapeutic strategy for preventing AG-induced hearing loss and provide a foundation for further mechanistic and translational investigations.

## METHODS

### Computational Scaffold-Hopping

To identify structurally related analogs with enhanced otoprotective activity, a computational scaffold-hopping strategy was employed using similarity-based virtual screening. Beginning with MR16728 as the initial query, a library of candidate analogs was generated through molecular fingerprint–based searches using the SpaceLight platform^18^.

Molecular similarity searches were performed using Connected Subgraph Fingerprints (CSFP), a topological fingerprinting approach that encodes molecular structures as exhaustive sets of connected subgraphs representing atomic environments and substructural features^19,20^. CSFP methods capture broader structural relationships, including functional group arrangements and connectivity patterns, enabling identification of analogs with conserved pharmacophores but modified scaffolds.

Computational Scaffold-Hopping was performed against the National Cancer Institute Developmental Therapeutics Program (NCI/DTP) compound repository, which contains more than 320,000 chemically diverse small molecule compounds available for screening and drug discovery^21^. Similarity between candidate molecules and query structures was quantified using the Tanimoto coefficient (Tc), a widely used metric for comparing molecular fingerprints^22^. A Tc cutoff of 0.5 was applied to balance structural similarity with chemical diversity, allowing sufficient flexibility for scaffold hopping while maintaining core features of the lead compounds. Compounds meeting this threshold were selected for experimental validation. This scaffold-hopping workflow enabled the identification of a focused library of 145 structurally related analogs for downstream *in vivo* screening, facilitating the discovery of compounds with improved otoprotective efficacy.

### Animals

Zebrafish (*Danio rerio*) embryos were obtained from in-house breeding of adult wild-type (AB strain) and transgenic lines maintained at the Idaho State University zebrafish facility. Fish were housed under standard laboratory conditions at 28.5 °C on a 14:10 h light/dark cycle in a recirculating aquatic system, in accordance with established zebrafish husbandry protocols^23,24^. Embryos were collected via natural spawning and raised in embryo medium (994 μM MgSO₄, 150 μM KH₂PO₄, 42 μM Na₂HPO₄, 986 μM CaCl₂, 503 μM KCl, 14.9 mM NaCl, and 714 μM NaHCO₃; pH 7.2) at 28 °C until 5–7 days post-fertilization (dpf), the developmental stage at which lateral line neuromast hair cells are fully functional and suitable for ototoxicity assays^25,26^.

Both wild-type and transgenic *Tg(brn3c:GFP)* fish were used depending on the experimental application, including dye uptake and confocal imaging assays. Fish were maintained in embryo medium (EM) and staged according to established morphological criteria^25^. Prior to experimental procedures, fish were screened to ensure normal development and the absence of morphological abnormalities.

All experimental procedures were conducted in compliance with institutional animal care and use guidelines and were approved by the Idaho State University Institutional Animal Care and Use Committee.

### Reagents

Aminoglycosides used in this study included gentamicin sulfate (Gold Biotechnology), neomycin sulfate, tobramycin, amikacin sulfate, and streptomycin sulfate (Millipore Sigma). All AGs were obtained as powders and dissolved in EM to prepare stock solutions, which were subsequently diluted to working concentrations immediately prior to use.

Candidate otoprotective compounds, including MR16728 and its structural analogs (e.g., 28510, 663865, and 27671), were obtained from the National Cancer Institute Developmental Therapeutics Program (NCI/DTP) as dry powders. Compounds were dissolved in dimethyl sulfoxide (DMSO; Sigma-Aldrich) to prepare 20 mM stock solutions, which were stored at −20°C. Compounds were diluted in EM to final working concentrations, ensuring that the final DMSO concentration did not exceed 1% (v/v), a level previously shown to have minimal effects on zebrafish viability and neuromast hair cell integrity^27^.

All reagents were prepared fresh or thawed immediately prior to use and handled according to manufacturer guidelines to ensure stability and reproducibility across experiments.

### Zebrafish Hair Cell Protection Assay

Hair cell protection assays were performed using larval zebrafish (5–7 dpf), a developmental stage at which lateral line neuromast hair cells are fully functional and highly sensitive to AG-induced ototoxicity^25,26^. Assays were conducted using established *in vivo* phenotypic screening approaches adapted for identifying otoprotective compounds^14,28^.

A whole fish neuromast hair cell scoring system was used to evaluate hair cell integrity. For each experimental condition, groups of 9–12 fish were placed in 24-well plates containing EM. Fish were pre-incubated with candidate compounds for 1 hour at either fixed concentrations (10, 25, and 50 μM) or across a broader range (0.01–100 μM) to determine the 50% hair cell protection concentration (HC_50_). Following pre-incubation, fish were co-incubated with AGs for protocol-specific durations.

Acute exposure assays consisted of 30-minute co-incubation with either 110 μM gentamicin or 50 μM neomycin, followed by complete media replacement with fresh EM and a 1-hour recovery period prior to assessment ^28^. Chronic assays involved 24-hour exposures in which fish underwent a 1-hour pre-incubation followed by 24-hour co-incubation with AGs prior to assessment^13,29^. AG concentrations used in the chronic assays included 200 μM gentamicin, 1 mM tobramycin, 10 mM amikacin, and 500 μM streptomycin. AG concentrations were determined through preliminary dose–response experiments to achieve approximately 80% neuromast hair cell loss, ensuring a robust yet submaximal ototoxic insult that permits detection of protective effects. To maintain consistent injury levels across experimental cohorts, AG doses were adjusted as needed.

### Lateral Line Neuromast Imaging and Scoring

Following treatment, fish were stained with 0.005% DASPEI (2-[4-(dimethylamino)styryl]-N-ethylpyridinium iodide; Sigma-Aldrich) for 15 minutes to label mitochondria within hair cells and associated neuronal structures. DASPEI is a well-established vital dye for assessing neuromast integrity and hair cell viability in zebrafish lateral line assays^28,30^. After staining, fish were rinsed with EM and anesthetized using 0.01% Syncaine (pharmaceutical grade tricaine methanesulfonate; Syndel) to immobilize specimens for imaging.

Fish were imaged using a Leica M165 fluorescence stereomicroscope under consistent illumination and exposure settings across all experimental groups. Neuromast hair cell integrity was assessed using a quantitative whole-fish scoring system based on fluorescence intensity and morphology. Neuromasts were evaluated on a scale from 0 to 4, where a score of 0 indicated complete loss of fluorescence and severe hair cell damage comparable to AG-treated controls, and a score of 4 indicated intact, bright fluorescence comparable to untreated controls. Intermediate scores reflected partial preservation of neuromast hair cell morphology and fluorescence (Figure S1).

For each fish, scores from multiple neuromasts distributed along the lateral line were averaged to generate a whole-fish neuromast score. Group-level outcomes were calculated as the mean ± SEM and normalized to untreated control groups and expressed as percentage of the mean neuromast score of untreated controls. This scoring approach enables rapid, reproducible, and quantitative assessment of hair cell damage and protection *in vivo*^12,14,30^.

### FM1-43 Uptake

To investigate whether candidate otoprotective compounds modulate AG entry through the MET channels, FM1-43 uptake assays were performed in larval zebrafish. The styryl dye FM1-43 (N-(3-triethylammoniumpropyl)-4-(4-(dibutylamino)styryl)pyridinium dibromide; Thermo Fisher Scientific) is a well-established probe that rapidly enters hair cells through open MET channels, and its intracellular fluorescence intensity serves as a proxy for MET channel permeability^31,32^.

Larval fish (5–7 dpf) were pre-incubated with candidate compounds for 1 hour at specified concentrations, followed by brief co-incubation with 3 μM FM1-43 for 45 seconds. This short exposure period preferentially labels hair cells via MET channel uptake while minimizing nonspecific dye internalization^32^. Following incubation, fish were rinsed once with EM to remove excess dye and anesthetized with 0.01% Syncaine for imaging.

Fluorescence imaging was performed using a Leica M165 stereomicroscope under consistent acquisition settings. High-resolution images of defined neuromasts (P1 and P4) were captured using a SPOT Insight 4.0 MP FireWire monochrome camera. Quantitative analysis of fluorescence intensity was conducted using ImageJ^33^. Regions of interest (ROIs) corresponding to individual neuromasts were manually selected, and mean fluorescence intensity values were calculated. For each larva, fluorescence values from P1 and P4 neuromasts were averaged, and group-level means were computed for comparison across treatment conditions.

### AG Entry

To directly assess AG entry into hair cells, uptake of Texas Red-conjugated gentamicin (GTTR; AAT Bioquest) was evaluated in transgenic fish. GTTR is a fluorescently labeled AG that enables visualization of drug accumulation within hair cells and has been widely used to study AG trafficking and uptake mechanisms^34,35^.

*Tg(brn3c:GFP)* transgenic fish expressing hair cell-specific fluorescence at 5–7 dpf were pre-incubated with candidate compounds for 1 hour at specified concentrations. Fish were then co-incubated with 25 μM GTTR for 30 seconds to allow rapid uptake into hair cells. Following exposure, fish were rinsed once with fresh EM to remove excess dye and anesthetized with 0.01% MS-222 for imaging.

Confocal imaging was performed using a Nikon C2+ Ti-U confocal microscope. Images were acquired using FITC and far-red channels to visualize GFP-labeled hair cells and GTTR fluorescence, respectively. Neuromasts were imaged under consistent acquisition settings across all treatment groups to enable quantitative comparison. This assay complements FM1-43 uptake studies by directly measuring AG accumulation within hair cells.

### AG Antibiotic Activity

To evaluate whether candidate otoprotective compounds interfere with the antibacterial efficacy of AGs, antibiotic susceptibility testing was performed using a standard Kirby–Bauer disk diffusion assay^36,37^. This assay enables quantitative assessment of bacterial growth inhibition and is widely used to evaluate potential drug–drug interactions affecting antibiotic activity.

Sterile paper disks were prepared with AGs alone, candidate compounds alone, and combinations of AGs with candidate compounds. AG control disks were prepared using standard doses (10 μg gentamicin and 30 μg neomycin), while experimental compound disks were loaded with amounts approximating concentrations used in zebrafish assays (equivalent to 50 μM upon diffusion). Combination disks contained both AGs and candidate compounds at the same respective doses. Additional controls included blank paper disks, 1% DMSO vehicle controls, and chloramphenicol disks as a positive control for bacterial inhibition.

*E. coli* (ATCC 25922) cultures were prepared from 2-3 isolated colonies inoculated into Mueller–Hinton broth and grown with agitation at 37 °C for 2–3 hours to mid-log phase. Bacterial suspensions were adjusted to an optical density (OD600) of 0.08–0.1, corresponding to approximately 1–2 × 10⁸ CFU/mL, and then diluted 1:10 prior to plating. Mueller–Hinton agar plates were uniformly inoculated using a sterile swab in multiple directions to ensure confluent bacterial growth. After a 15-minute equilibration period, prepared disks were applied to the agar surface in labeled positions.

Plates were incubated at 35–37 °C for 18–24 hours, after which zones of inhibition were measured to the nearest millimeter. Using inhibition zone diameters, areas of inhibition (mm^2^) were calculated for each disk, and an average zone of inhibition was calculated for each experimental group. Comparisons of inhibition zone diameters between AG-only and AG-plus-compound conditions were used to determine whether candidate compounds altered AG antibacterial activity. This approach provides a robust and standardized method for assessing potential antagonistic or synergistic interactions^36,37^.

### ADME Prediction

To evaluate the drug-like properties and translational potential of candidate compounds, *in silico* ADME predictions were performed using the QikProp module^38^. QikProp is a widely used computational tool that estimates key pharmacokinetic descriptors based on physicochemical properties and established quantitative structure–property relationship (QSPR) models^39^.

For each compound, the following parameters were calculated: molecular weight, predicted Caco-2 cell permeability (QPPCaco), blood–brain barrier partition coefficient (QPlogBB), and human oral absorption (HOA). These descriptors were selected to assess critical aspects of drug-likeness, including oral absorption, tissue distribution, and potential cardiotoxicity risk. Predicted values were interpreted relative to reference ranges derived from known drug compounds, as implemented in QikProp.

Chemical structures were prepared using standard ligand preparation workflows prior to ADME prediction to ensure appropriate protonation states and geometry optimization. ORC-13661, a clinically advancing otoprotective compound, was included as a reference comparator to contextualize the predicted pharmacokinetic properties of the candidate analogs^12,13^. These *in silico* assessments provided an initial framework for evaluating the developability of lead compounds and informed subsequent optimization strategies.

## RESULTS

### Computational Scaffold-Hopping Yielded a Focused Compound Library

To identify molecular candidates with improved otoprotective efficacy across both acute and chronic AG treatments, a computational scaffold-hopping strategy was employed. Building on the observed acute otoprotective activity of MR16728, an ion channel modulator, we sought to identify compounds that retained acute efficacy while enhancing chronic protective performance.

A focused library of 145 structurally related compounds was generated based on the core scaffold of MR16728 using a similarity-based search strategy employing Connected Subgraph Fingerprints (CSFP). CSFP encodes molecular features such as atomic environments and substructural connectivity. Structural similarity between compounds was quantified using the Tanimoto coefficient (Tc), a widely used metric in cheminformatics for comparing molecular fingerprints and identifying analogs. A Tc cutoff of 0.5 was applied to balance sufficient structural diversity with retention of key scaffold features of the parent compound.

### Initial Screen Against Acute Gentamicin Exposure

Initial *in vivo* zebrafish screening identified multiple analogs with enhanced acute otoprotective activity relative to MR16728. The structures of the top candidate compounds (28510, 663865, and 27671) are shown in Figure 1. These three compounds emerged as top candidates based on initial neuromast hair cell studies that showed their robust protection (at 25 μM) against 110 μM Gentamicin, an AG-treated control condition that reliably produced approximately 80–90% hair cell damage (Figure 2). In these studies, the three top candidate compounds restored the %neuromast score to >80%. These top compounds were subsequently prioritized for further dose–response characterization and evaluation across acute and chronic AG exposure assays (Figures 3 and 4).

**Figure 1.**
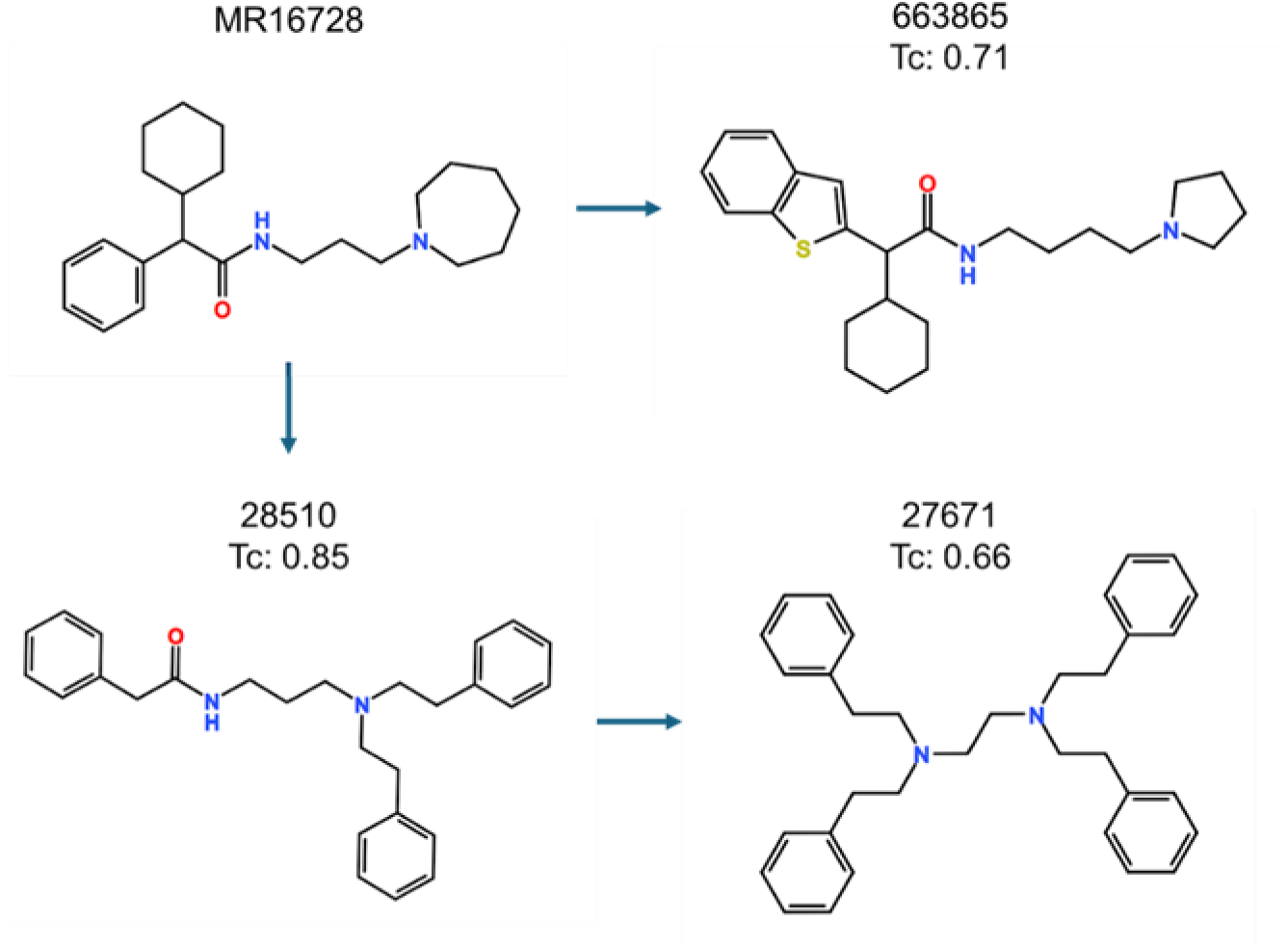
Chemical structures for MR16728 and the top three MR16728-derived analogs with corresponding Tanimoto similarity coefficients (Tc).

**Figure 2.**
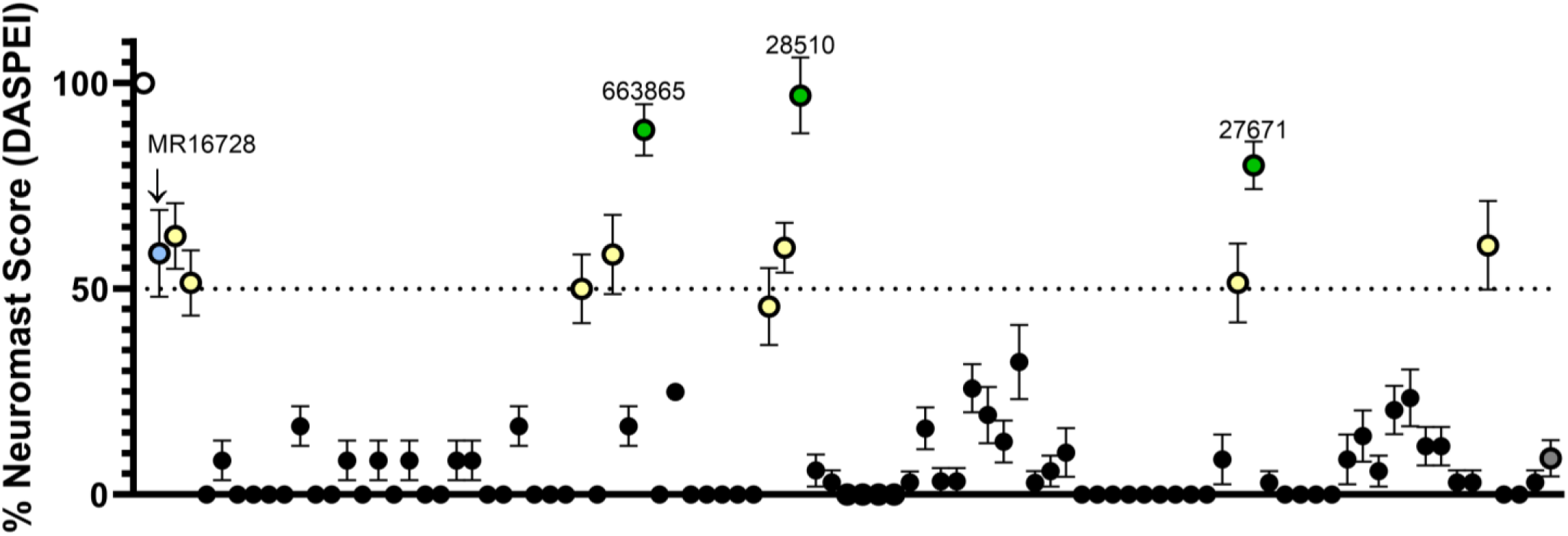
Initial screening of MR16728 and MR16728-derived analogs at 25 μM for protection against acute gentamicin exposure (110 μM) in zebrafish. The white point (far left) indicates untreated controls, while the dark grey point (far right) indicates 110 μM gentamicin-treated controls. The blue point represents MR16728, the yellow points represent several compounds that revealed activity similar to that of MR16728, and the green points represent the 3 compounds of interest that reached >80% neuromast hair cell restoration. Values represent mean whole-fish neuromast quality expressed as a percentage of untreated controls ± SEM (*n* = 6).

**Figure 3.**
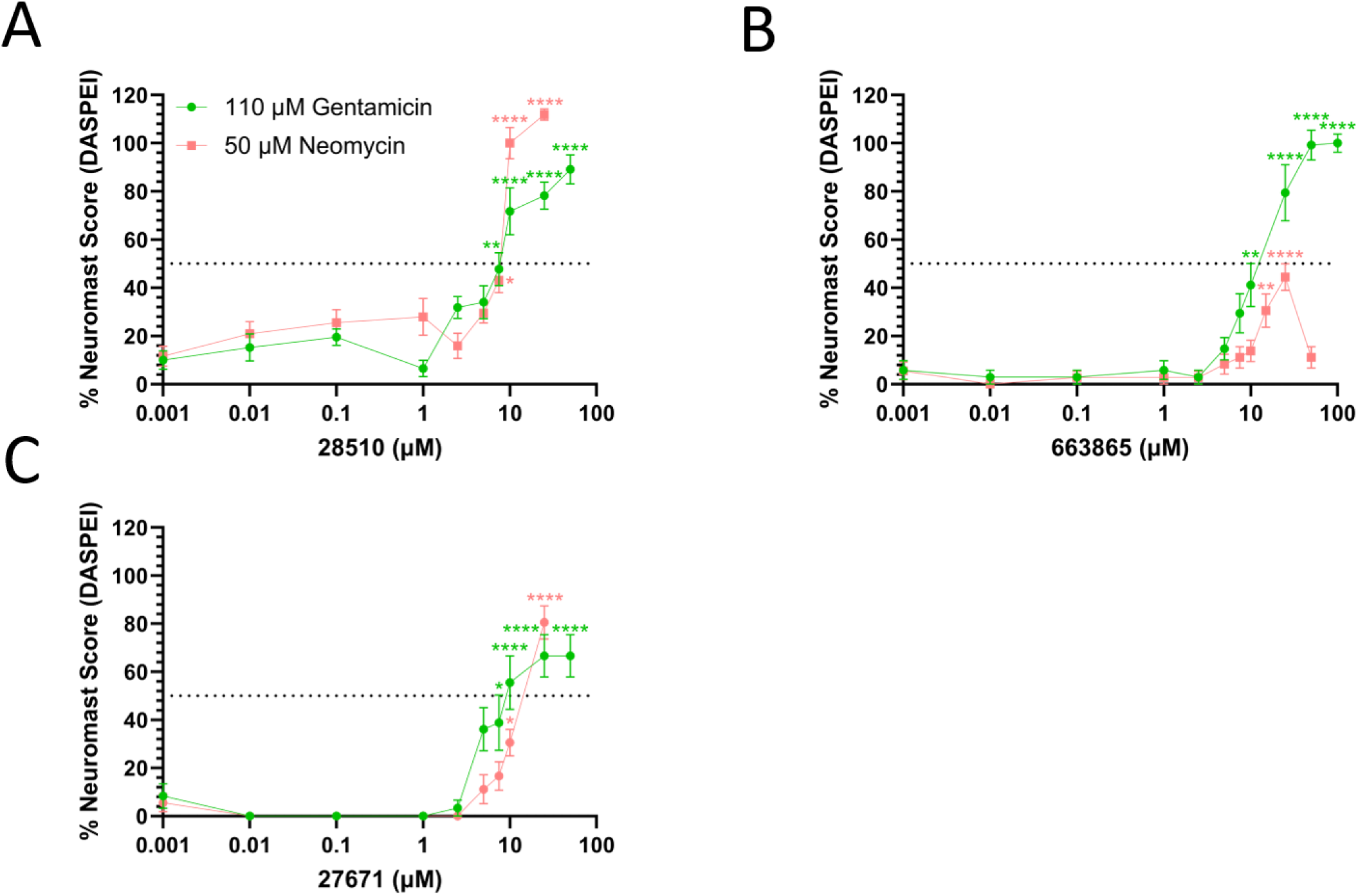
Hit compound protection against acute AG-induced hair cell death following pre-exposure to A) 28510, B) 663865, and C) 27671. Fish were challenged with 110 μM gentamicin or 50 μM neomycin. Values represent mean whole-fish neuromast hair cell quality expressed as a percentage of untreated controls ± SEM (*n* = 9-12). Statistical significance was determined by one-way ANOVA with Tukey’s post hoc test (*p<0.05, **p<0.01, ***p<0.001, ****p<0.0001), with asterisks indicating comparisons to the AG-treated controls. A dose of 0.001 represents AG-only treatment. HC_50_ values were estimated by linear regression between concentrations above and below 50% neuromast hair cell quality.

**Figure 4.**
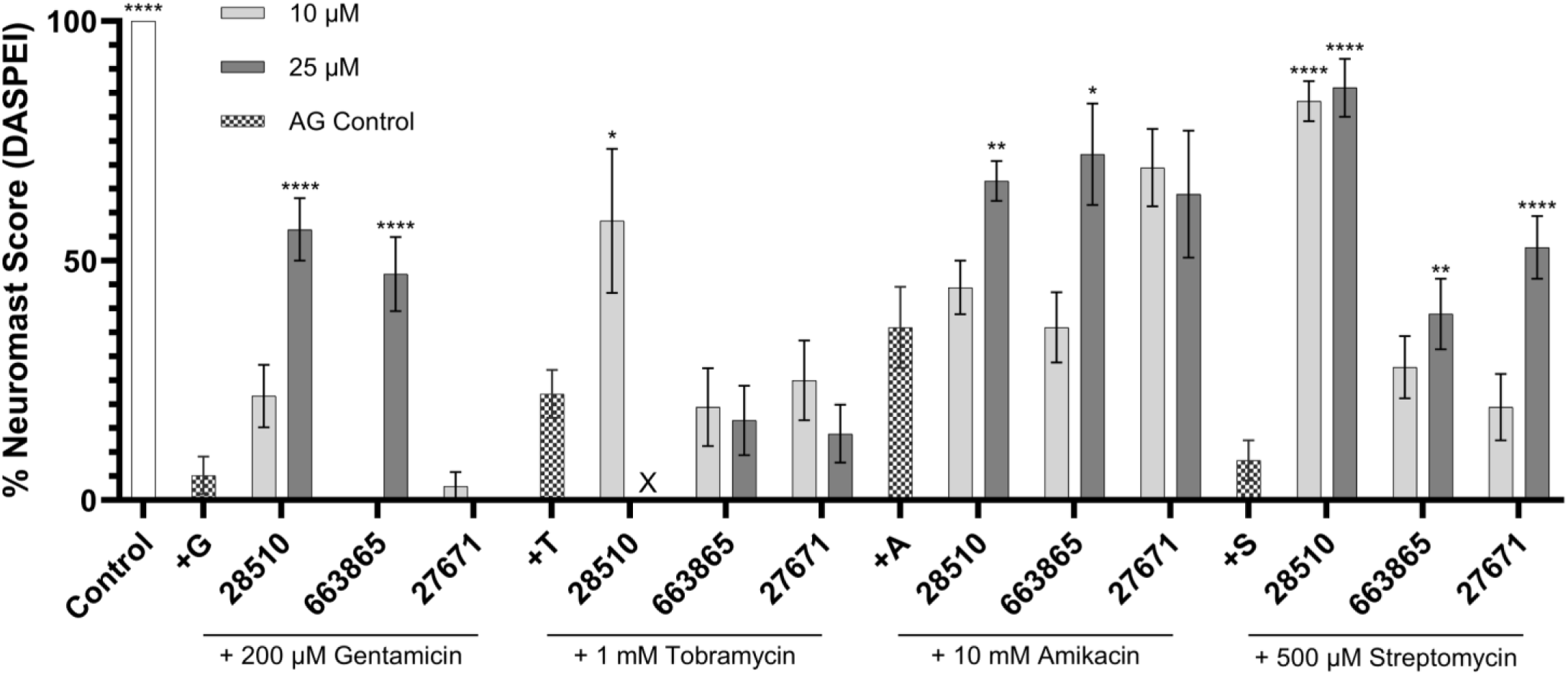
Protection against chronic AG exposures. Values represent mean whole-fish neuromast hair cell quality expressed as a percentage of untreated controls (indicated by the white bar on the far left) ± SEM (*n* = 9). Statistical significance was determined by one-way ANOVA with Tukey’s post hoc test (*p<0.05, **p<0.01, ***p<0.001, ****p<0.0001). Black asterisks indicate comparisons versus corresponding AG-treated controls (+G = gentamicin, +T = tobramycin, +A = amikacin, +S = streptomycin). Absence of data indicates neuromast hair cell quality scores of 0%, while an X indicates total fish mortality.

### Protection Against Acute AG Exposure

The top candidate compounds (28510, 663865, and 27671) were evaluated for their ability to protect zebrafish lateral line neuromast hair cells from acute AG exposure against 110 μM gentamicin or 50 μM neomycin using established protocols that reliably induce substantial hair cell loss^14,28^.

Compound 28510 demonstrated the most robust and consistent otoprotective activity. Against gentamicin, significant protection was observed at 7.5 μM, with protection continuing to increase and reaching maximal protection of 90% at 50 μM (p<0.0001, Figure 3A). The calculated HC_50_ value for gentamicin protection was 7.74 μM. Similarly, 28510 provided significant protection against neomycin-induced damage starting at 7.5 μM, achieving 100% protection at 10 μM (p<0.0001), with an HC_50_ of 7.80 μM (Figure 3A).

Compound 663865 provided significant protection beginning at 10 μM with protection continuing to increase and reaching maximal protection of 100% at 25 μM (p<0.0001, Figure 3B). The calculated HC_50_ value for gentamicin protection was 13.46 μM. In contrast, protection against neomycin was limited. Neuromast integrity was not restored to the untreated control levels. An HC_50_ value could not be determined despite showing significant improvement in neuromast quality at 25 μM relative to AG-treated controls (Figure 3B).

Compound 27671 displayed significant protection against gentamicin beginning at 7.5 μM with a maximal protection of 66% at 25 μM (p<0.0001). An HC_50_ of 9.17 μM was determined. Significant protection against neomycin was observed at 10 μM, with maximal protection of 80% at 25 μM (p<0.0001). An HC_50_ of 15.83 μM was determined (Figure 3C).

These results demonstrate that all three compounds confer protection against AG toxicity, with 28510 showing the greatest potency and efficacy across both gentamicin and neomycin.

### Protection Against Chronic AG Exposure

The three lead compounds (28510, 663865, and 27671) were also evaluated for their ability to protect against chronic AG-induced hair cell death (Figure 4), which better represents clinically relevant drug exposure conditions^14,28,29^. Initial screens were performed at 10 and 25 μM across multiple AGs, including gentamicin, tobramycin, amikacin, and streptomycin. Full dose–response analyses were subsequently conducted for compounds demonstrating promising activity.

Compound 28510 demonstrated broader protection against multiple AGs. Significant improvements relative to AG-treated controls were observed for gentamicin and streptomycin (p<0.0001), with partial protection also evident against tobramycin and amikacin. Compound 663865 significantly protected hair cells from gentamicin-induced loss (p<0.0001), while showing only partial protection against amikacin and streptomycin. Compound 27671 displayed minimal activity overall, with significant improvement observed against streptomycin-treated at 50 μM (p<0.0001) (Figure 4).

These findings demonstrate that 28510 provides the most robust and broad-spectrum protection under both acute and chronic exposure conditions (summarized in Table 1), supporting its selection as the top candidate for subsequent mechanistic and translational studies.

**Table 1.** Summary of protective effects of lead compounds across all AG exposure assays.

|  | Statistical Significance of Neuromast Hair Cell Protection<br>Relative to AG-Treated Controls |  |  |  |  |  |
| --- | --- | --- | --- | --- | --- | --- |
| Compound | 28510 |  | 663865 |  | 27671 |  |
| Acute Exposure | 25 µM | 50 µM | 25 µM | 50 µM | 25 µM | 50 µM |
| 110 µM Gentamicin | **** | **** | **** | **** | **** | **** |
| 50 µM Neomycin | **** | **** | **** | ns | **** | **** |
| Chronic Exposure | 10 µM | 25 µM | 10 µM | 25 µM | 10 µM | 25 µM |
| 200 µM Gentamicin | ns | **** | ns | **** | ns | ns |
| 1 mM Tobramycin | * | X | ns | ns | ns | ns |
| 10 mM Amikacin | ns | ** | ns | * | ns | ns |
| 500 µM Streptomycin | **** | **** | ns | ** | ns | **** |
| Values represent mean whole-fish neuromast quality expressed as a percentage of untreated controls ± SEM (*n* = 9). Statistical significance was determined by one-way ANOVA with Tukey's post hoc test (* *p* <0.05, ** *p* <0.01, *** *p* <0.001, **** *p* <0.0001) versus AG-treated controls. X = zebrafish mortality, ns = not significant. |  |  |  |  |  |  |

### Focused Evaluation of Top Lead Compound 28510

Following initial screening, detailed dose–response analyses were performed to further characterize the chronic otoprotective effects of compound 28510 and to determine HC_50_ values across clinically relevant AGs, including gentamicin, tobramycin, amikacin, and streptomycin^13^.

In the chronic AG exposure assays, 28510 exhibited markedly enhanced protective activity. Against high dose 200 μM gentamicin, there was a maximal recovery of 43% at 50 μM. While this failed to achieve a response of at least 50%, the effect was a statistically significant improvement in neuromast hair cell quality (p<0.0001) from gentamicin-treated controls. When gentamicin doses were reduced to 25 μM, potent protection was observed at 25 μM with maximal recovery reaching 100% of untreated control. A low HC_50_ of 4.31 μM was determined. Similarly, in tobramycin exposure assays, 10 μM 28510 achieved 80% neuromast hair cell recovery compared to untreated controls. The HC_50_ was determined to be 6.25 μM. Protection against amikacin was moderate.

Although a maximal response of 62% was observed at 25 μM (HC_50_ = 8.75 μM), this protective effect was not statistically significant relative to amikacin-treated controls. Against streptomycin, 28510 achieved a maximal response of 75% at 50 μM, with an HC_50_ of 14.37 μM, representing statistically significant protection relative to streptomycin-treated controls (p<0.0001, Figure 5).

**Figure 5.**
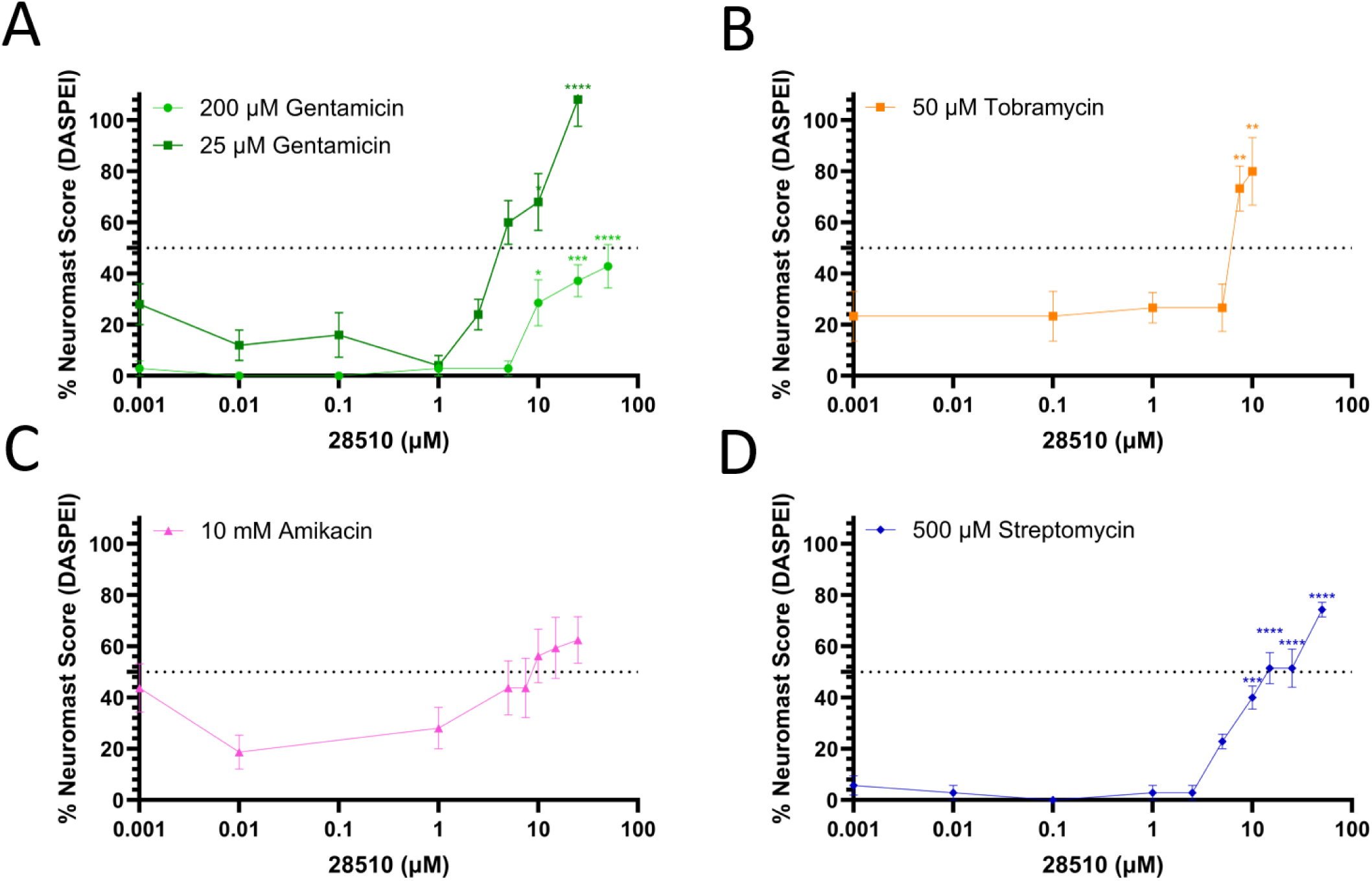
Lead compound 28510 dose-response against chronic AG exposures. Values represent mean whole-fish neuromast hair cell quality expressed as a percentage of untreated controls ± SEM (*n* = 9). Statistical significance was determined by one-way ANOVA with Tukey’s post hoc test (*p<0.05, **p<0.01, ***p<0.001, ****p<0.0001) versus AG-treated controls. A dose of 0.001 μM represents AG-only treatment. HC_50_ values were estimated by linear regression between concentrations above and below 50% neuromast hair cell quality. Data points in red indicate zebrafish mortality.

These detailed dose-response results further demonstrate that 28510 exhibits potent, dose-dependent, and broad-spectrum protection against AG-induced ototoxicity under chronic exposure conditions, supporting its potential as a lead compound for mitigating AG-induced ototoxicity.

### MET Channel Activity – FM1-43 Uptake

To determine whether the otoprotective effects of the lead compounds were mediated through inhibition of MET channel activity, FM1-43 uptake assays were performed in zebrafish lateral line hair cells. A reduction in FM1-43 fluorescence relative to untreated controls was interpreted as evidence of MET channel inhibition, whereas unchanged fluorescence indicated that compound-mediated otoprotection occurred independently of MET channel blockade.

Pre-incubation with compounds 28510, 663865, and 27671 did not significantly reduce FM1-43 uptake compared to untreated controls. In all cases, fluorescence levels remained comparable to control groups, indicating that these compounds do not inhibit MET channel-mediated dye entry (Figure 6). In contrast, EDTA-treat neuromasts, a positive control for MET channel disruption, exhibited significantly reduced fluorescence, consistent with impaired channel function.

**Figure 6.**
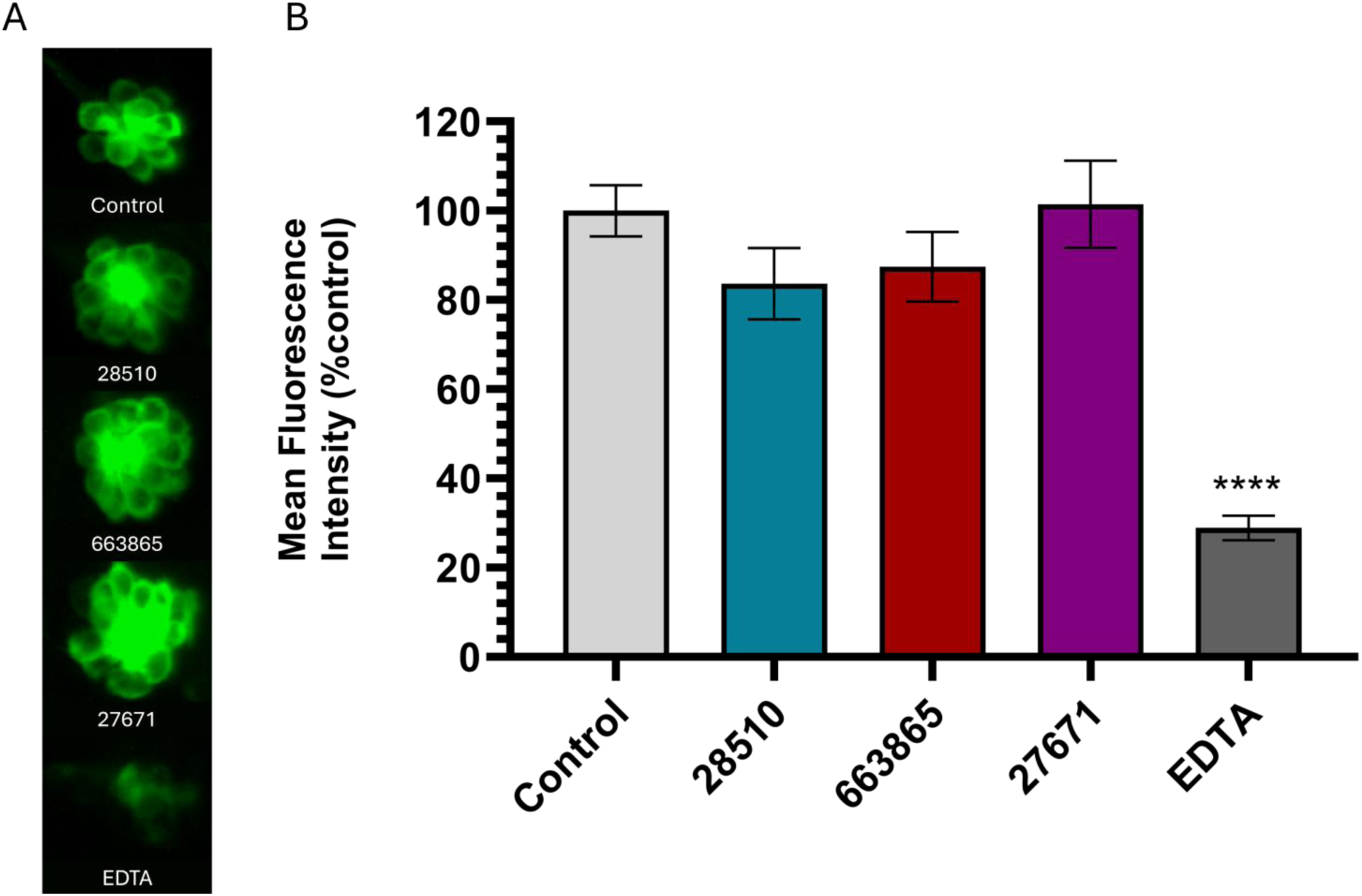
A) FM1-43 uptake in wild-type zebrafish neuromasts following 1-hour exposure to 50 μM 28510, 663865, 27671, and EDTA. B) Average fluorescent intensity of FM1-43 dye uptake as represented as a percentage of untreated controls ± SEM (*n* = 12-15). Statistical significance was determined by one-way ANOVA with Tukey’s post hoc test (*p<0.05, **p<0.01, ***p<0.001, ****p<0.0001) versus untreated control.

Furthermore, fluorescence intensity in compound-treated groups was significantly greater than that observed in EDTA-treated controls, reinforcing the conclusion that MET channel activity remained intact in the presence of these compounds.

### MET Channel Activity – AG Entry

To directly evaluate whether candidate compounds affect AG uptake into hair cells, GTTR loading assays were performed in zebrafish lateral line neuromasts. GTTR fluorescence provides a direct measure of AG entry into hair cells and has been widely used to assess drug uptake mechanisms^34^. A marked reduction or absence of the GTTR signal relative to untreated controls was interpreted as evidence of inhibited AG entry, consistent with MET channel blockade. In contrast, comparable levels of GTTR fluorescence indicated that compound-mediated otoprotection occurred independently of AG uptake inhibition.

Pre-incubation with 50 μM compounds 28510, 663865, and 27671 for 1 hour did not result in a significant reduction in GTTR fluorescence compared to untreated controls, indicating that these compounds do not inhibit AG entry into hair cells (Figure 7). Fluorescence intensity remained comparable across all compound-treated and control groups, suggesting that AG uptake pathways remain functionally intact. In contrast, berbamine-treated neuromasts, a positive control for inhibition of AG uptake, exhibited markedly reduced GTTR fluorescence.

**Figure 7.**
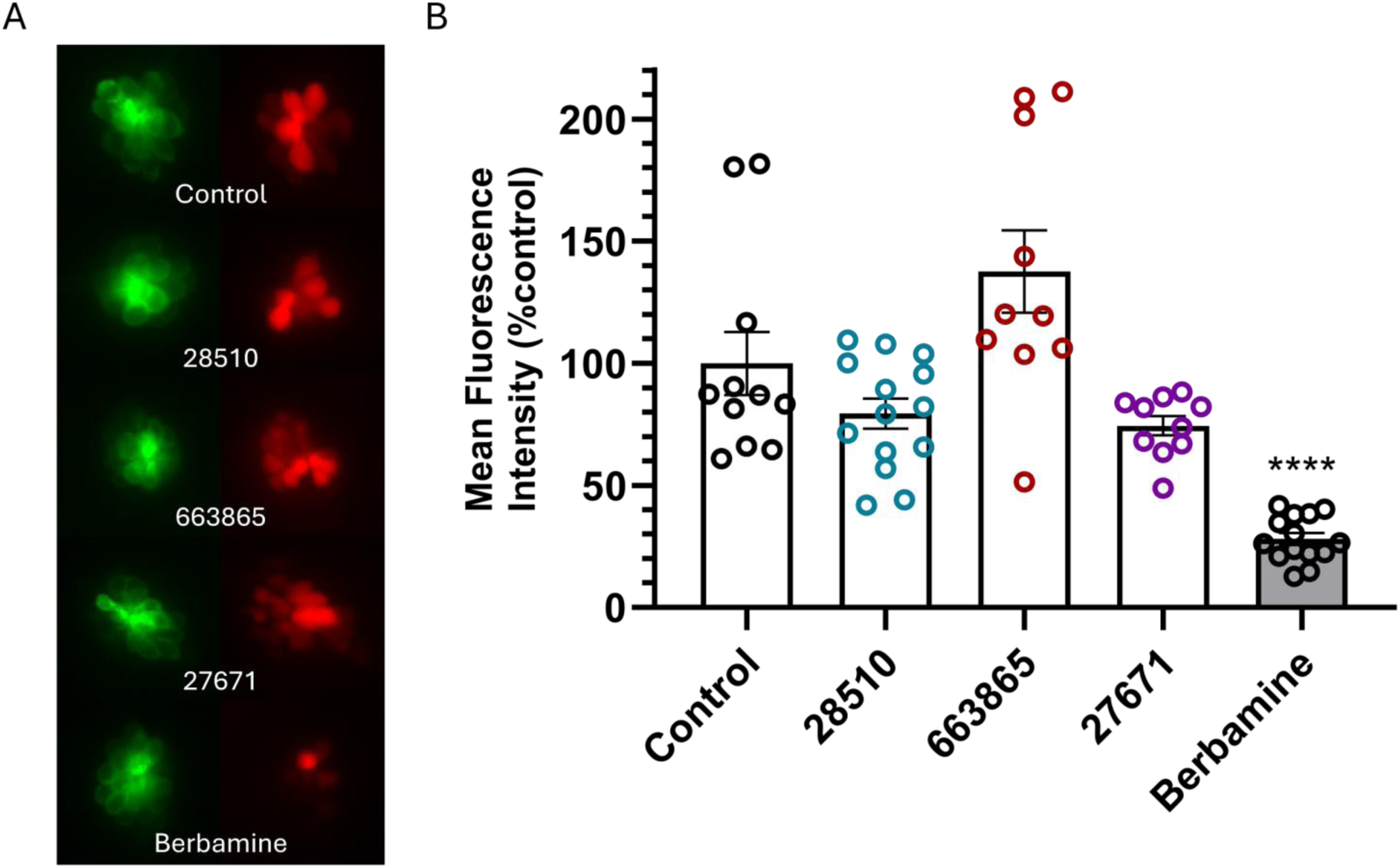
A) GTTR uptake in *Tg(brn3c:GFP)* lateral line neuromasts following exposure to 50 μM compounds 28510, 663865, 27671, and berbamine. B) Average fluorescent intensity of GTTR dye uptake as represented as a percentage of untreated controls ± SEM (*n* = 10-14). Statistical significance was determined by one-way ANOVA with Tukey’s post hoc test (*p<0.05, **p<0.01, ***p<0.001, ****p<0.0001) versus untreated control.

Together with the FM1-43 uptake results, these findings demonstrate that the otoprotective effects of 28510 and related analogs are not mediated through inhibition of MET channels. Instead, these compounds likely act through intracellular or downstream pathways that mitigate AG-induced hair cell damage.

### AG Antibiotic Activity

To evaluate whether the lead compound 28510 interferes with the antibacterial efficacy of AGs, disk diffusion assays were performed using *E. coli* (ATCC 25922), a standard reference strain for antimicrobial susceptibility testing. This analysis is critical for assessing the translational compatibility of candidate otoprotective agents with concurrent antibiotic therapy.

Co-administration of 28510 with either gentamicin or neomycin did not significantly alter zones of bacterial growth inhibition compared to AG treatment alone (Figure 8). Quantitative measurements of inhibition areas revealed no statistically significant differences between AG-only and AG+28510 conditions, indicating that 28510 does not compromise the bactericidal activity of these antibiotics.

**Figure 8.**
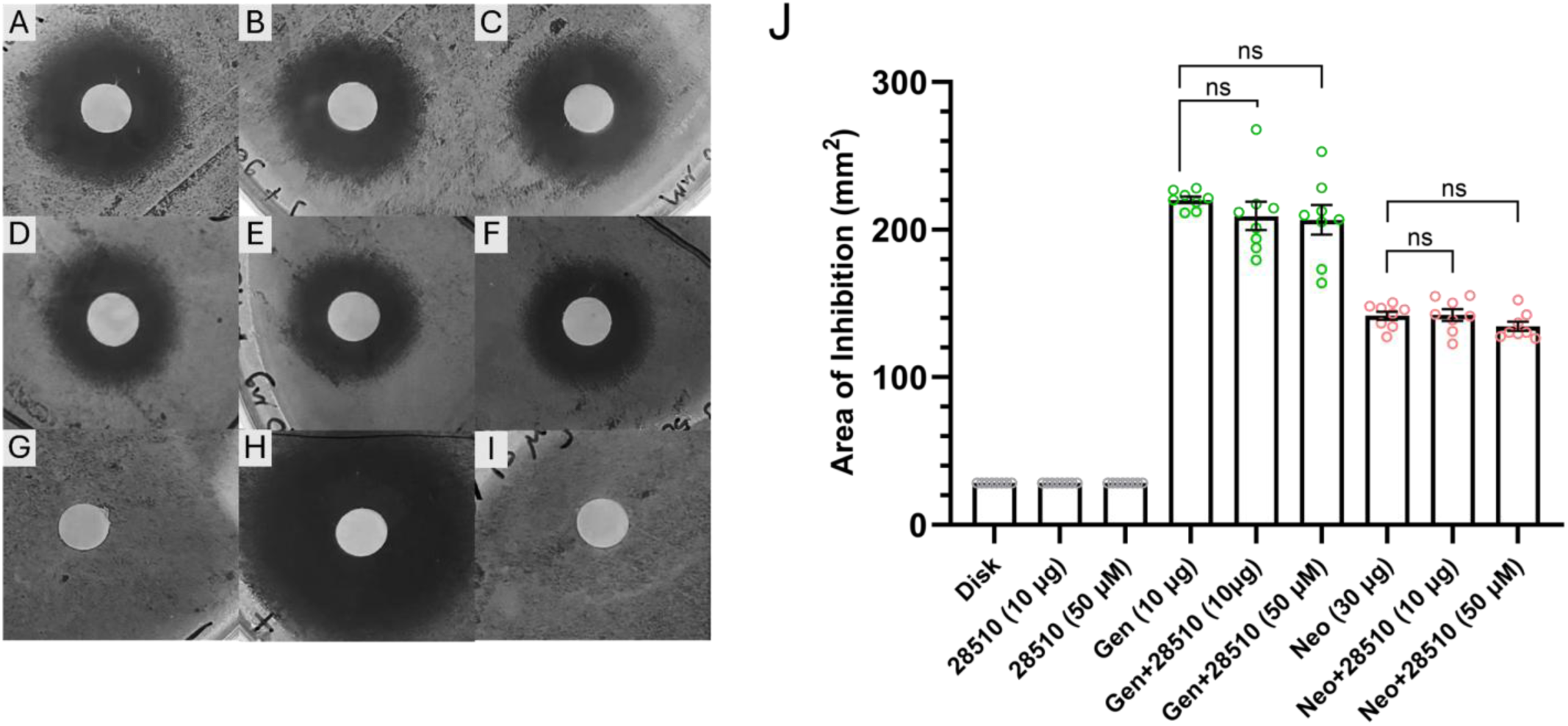
Comparison of *E. coli* growth inhibition zones across treatment conditions: A) gentamicin, B) 28510 (10 μg) + gentamicin, C) 28510 (50 μM) + gentamicin, D) neomycin, E) 28510 (10 μg) + neomycin, F) 28510 (50 μM) + neomycin, G) untreated disk, H) chloramphenicol, I) 28510 alone, and J) representative Kirby–Bauer disk diffusion assay using *E. coli* ATCC 25922. This assay evaluates the effect of 28510 on the bactericidal activity of gentamicin and neomycin. Values represent mean inhibition area (mm²) ± SEM (n = 8). Statistical significance was determined by one-way ANOVA with Tukey’s post hoc test (*p<0.05, **p<0.01, ***p<0.001, ****p<0.0001) versus AG-treated controls. Control disks measure 6 mm in diameter.

In addition, treatment with 28510 alone produced no measurable zones of inhibition, demonstrating that the compound lacks intrinsic antibacterial activity under the conditions tested. These findings indicate that 28510 preserves the antimicrobial efficacy of AGs while providing otoprotective effects, supporting its potential utility as a co-therapeutic agent.

### Predicted ADME Properties

All compounds fell within the acceptable molecular weight range for drug-like molecules (130–725 Da), supporting their suitability for further development. Predicted Caco-2 permeability values (QPPCaco) for all compounds exceeded 500, indicating a high likelihood of efficient intestinal absorption. Similarly, QPlogBB values for all compounds were within the acceptable range (−3 to 1.2), suggesting potential to cross the blood–brain barrier (BBB) or other tight epithelial barriers such as blood–labyrinth barrier (BLB). Although no established *in silico* ADME model currently exists for predicting permeability across the BLB, BBB permeability metrics such as QPlogBB are commonly used as an approximate surrogate due to the structural and physiological similarities between these barrier systems. Accordingly, the favorable QPlogBB values observed for these compounds suggest that they are likely BLB-permeable, supporting their potential to reach inner ear tissues *in vivo*. Predicted human oral absorption (HOA) was high (score = 3) for all compounds except 27671, which exhibited low predicted oral absorption (score = 1), indicating potential limitations in bioavailability.

Overall, these *in silico* analyses indicate that the identified compounds possess favorable pharmacokinetic properties, particularly with respect to absorption, tissue distribution, and potential inner ear accessibility (Table 2).

**Table 2.**
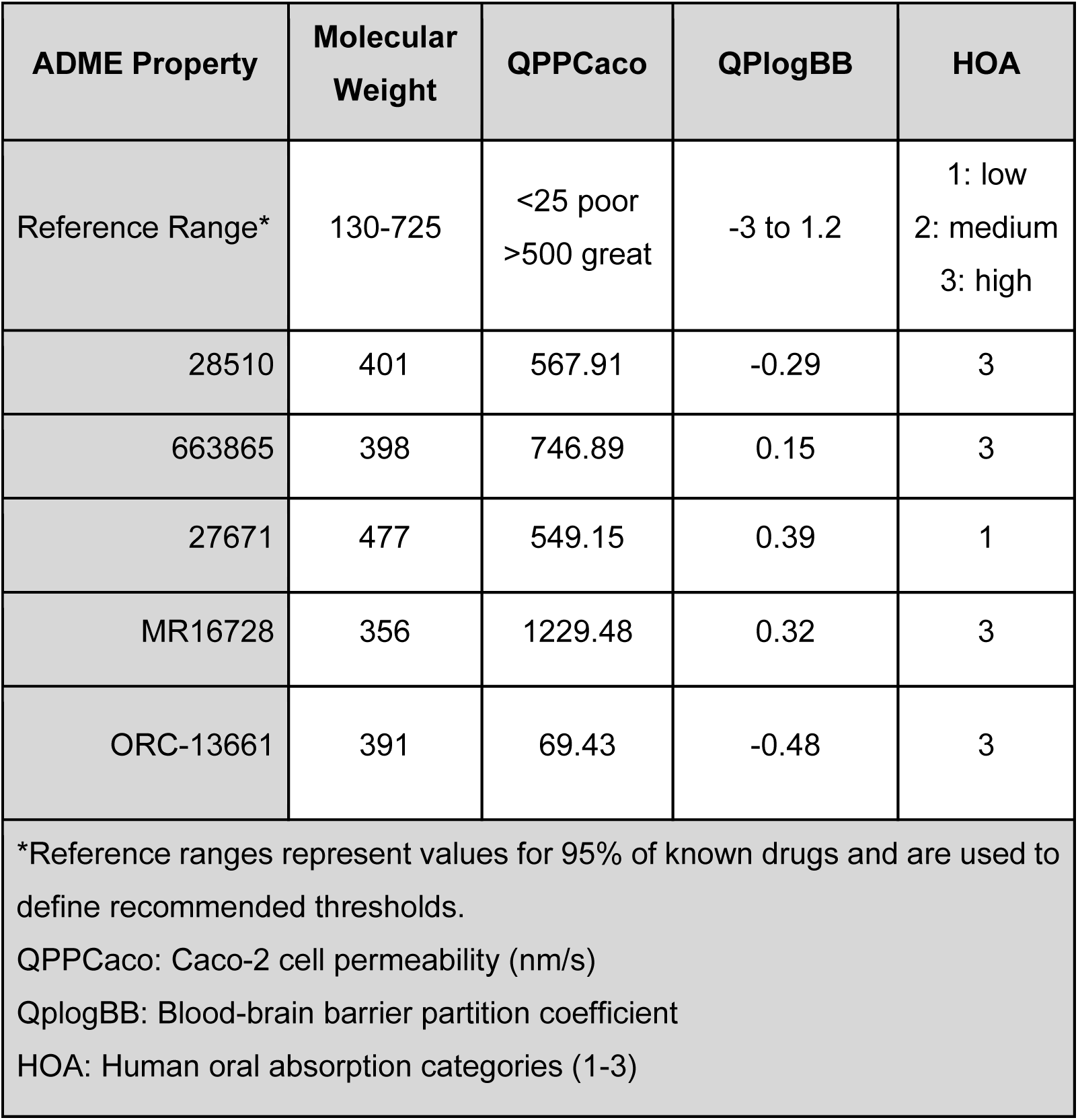
Predicted ADME properties.

| ADME Property | Molecular Weight | QPPCaco | QPlogBB | HOA |
| --- | --- | --- | --- | --- |
| Reference Range* | 130-725 | <25 poor<br>>500 great | -3 to 1.2 | 1: low<br>2: medium<br>3: high |
| 28510 | 401 | 567.91 | -0.29 | 3 |
| 663865 | 398 | 746.89 | 0.15 | 3 |
| 27671 | 477 | 549.15 | 0.39 | 1 |
| MR16728 | 356 | 1229.48 | 0.32 | 3 |
| ORC-13661 | 391 | 69.43 | -0.48 | 3 |
| *Reference ranges represent values for 95% of known drugs and are used to define recommended thresholds.<br>QPPCaco: Caco-2 cell permeability (nm/s)<br>QplogBB: Blood-brain barrier partition coefficient<br>HOA: Human oral absorption categories (1-3) |  |  |  |  |

## DISCUSSION

This study sought to identify novel compounds that mitigate AG-induced ototoxicity using a combined *in vivo* phenotypic screening and *in silico* scaffold-hopping approach. Building upon the previously identified ion channel modulator MR16728, which exhibits partial otoprotective effects but is limited by incomplete efficacy and dose-dependent toxicity^17^, we identified structurally related analogs with improved functional profiles. Among these, compound 28510 emerged as the most promising lead, demonstrating robust and broad-spectrum protection against multiple clinically relevant AGs, including gentamicin, tobramycin, amikacin, and streptomycin, across both acute and chronic exposures, highlighting the translational potential of this compound.

A key finding of this study is that 28510 achieves strong otoprotective efficacy at low micromolar concentrations while maintaining a favorable therapeutic window. Specifically, the effective concentrations required for protection (HC_50_ values in the low micromolar range) were consistently lower than concentrations associated with neuromast toxicity in zebrafish, indicating a clear separation between therapeutic and toxic dose ranges (Figure S2). This profile is consistent with other leading clinical candidates, such as ORC-13661, supporting the feasibility of dose optimization for translational development.

Mechanistically, our findings suggest that 28510 and related analogs do not exert their protective effects by inhibiting MET channel-mediated AG entry. Both FM1-43 and GTTR uptake assays demonstrated that these compounds do not significantly reduce dye or AG entry into hair cells, indicating that their protective activity likely occurs downstream of drug uptake. This distinct, non-MET-mediated action sets 28510 apart from leading clinical candidates such as ORC-13661, which function as MET channel blockers but do not address alternative AG uptake pathways, including endocytosis and permeation through transient receptor potential (TRP) channels^8^. Instead, the protective effects of 28510 likely arise from modulation of intracellular pathways downstream of AG entry. Given the established roles of calcium dysregulation, mitochondrial dysfunction, and ROS generation in AG-induced hair cell death, it is plausible that 28510 acts by stabilizing intracellular calcium homeostasis, preserving mitochondrial integrity, or attenuating oxidative stress signaling. These hypotheses warrant further investigation using targeted functional assays and transgenic models.

Combination strategies were also explored to assess potential synergistic effects. However, combining lead compounds did not yield meaningful improvements in hair cell protection. The 28510+663865 combination modestly increased tolerability at higher cumulative doses but did not enhance protective efficacy beyond that of 28510 alone. Similarly, while the 27671+663865 combination showed limited improvements under specific conditions, it also introduced increased toxicity and failed to demonstrate consistent or robust synergy (Figures S3–S5). Overall, these findings indicate that these compounds may share convergent downstream protective pathways, and that 28510 alone provides near-maximal efficacy within its class. Consequently, combinations of compounds from the same class do not appear to offer an advantage over monotherapy with the lead compound.

An important translational consideration is whether otoprotective agents interfere with the antimicrobial efficacy of AGs. In this study, 28510 did not alter the bactericidal activity of gentamicin or neomycin in standard disk diffusion assays, indicating compatibility with concurrent antibiotic therapy. This represents a critical advantage, as many candidate otoprotectants fail due to antagonistic drug–drug interactions.

From a medicinal chemistry perspective, the scaffold-hopping approach successfully identified structurally related analogs with improved biological activity, highlighting the utility of similarity-based virtual screening in phenotypic drug discovery. Preliminary structure–activity relationship insights suggest that conserved features, including amide linkages, hydrophobic substituents, and nitrogen-containing functional groups, may contribute to activity. In parallel, *in silico* ADME predictions indicate that 28510 possesses favorable pharmacokinetic properties, including high predicted intestinal and oral absorption as well as permeability across the gut-blood barrier, blood-brain barrier (BBB), and potentially the blood-labyrinth barrier (BLB), comparable to the clinically advancing otoprotective candidate ORC-13661. Further mechanistic studies and medicinal chemistry optimization will be essential to advance compound 28510 toward clinical application.

In summary, this study identifies 28510 as a promising first-in-class otoprotective lead that functions independently of MET channel blockade. Its broad-spectrum activity across multiple AGs and acute/chronic exposures, combined with preservation of AG antibiotic efficacy and a favorable therapeutic window, underscores its therapeutic potential. Moreover, the identification of a non-MET-mediated mechanism of protection opens new avenues for combination therapies, where intracellular protective agents like 28510 could be used alongside MET channel blockers to achieve synergistic protection against AG-induced hearing loss.

## STATEMENTS

### Ethics Statement

The animal study was approved by the Idaho State University Institutional Animal Care and Use Committee. The study was conducted in accordance with the legislative and institutional requirements.

### Author Contributions

EK: Conceptualization, Formal analysis, Investigation, Methodology, Validation, Visualization, Writing – original draft, Writing – review & editing.

CN: Formal analysis, Investigation, Methodology, Software, Visualization, Writing – review & editing.

SER: Conceptualization, Methodology, Facility management, Writing – review & editing.

TH: Investigation, Methodology, Writing – review & editing.

CL: Investigation, Methodology, Visualization, Writing – review & editing.

KF: Investigation, Methodology, Writing – review & editing.

KAC: Conceptualization, Formal analysis, Supervision, Writing – review & editing.

DX: Conceptualization, Funding acquisition, Project administration, Supervision, Data curation, Writing – review & editing.

## Acknowledgement

We thank the National Cancer Institute Developmental Therapeutics Program (NCI/DTP) https://dtp.cancer.gov for providing compounds present in this publication. Specifically, NSC#: 28510, 663865, 27671, 662128, etc. We thank Dr. David Raible and Patricia Wu from the Raible Lab at the University of Washington School of Medicine for generously providing the *Tg(brn3c:GFP)* transgenic zebrafish line, which was essential for imaging and mechanistic studies in this work.

