## Supplemental Materials for "Discovery of a Novel Non-MET-Mediated Otoprotective Compound Against Aminoglycoside-Induced Ototoxicity"

### TITLE

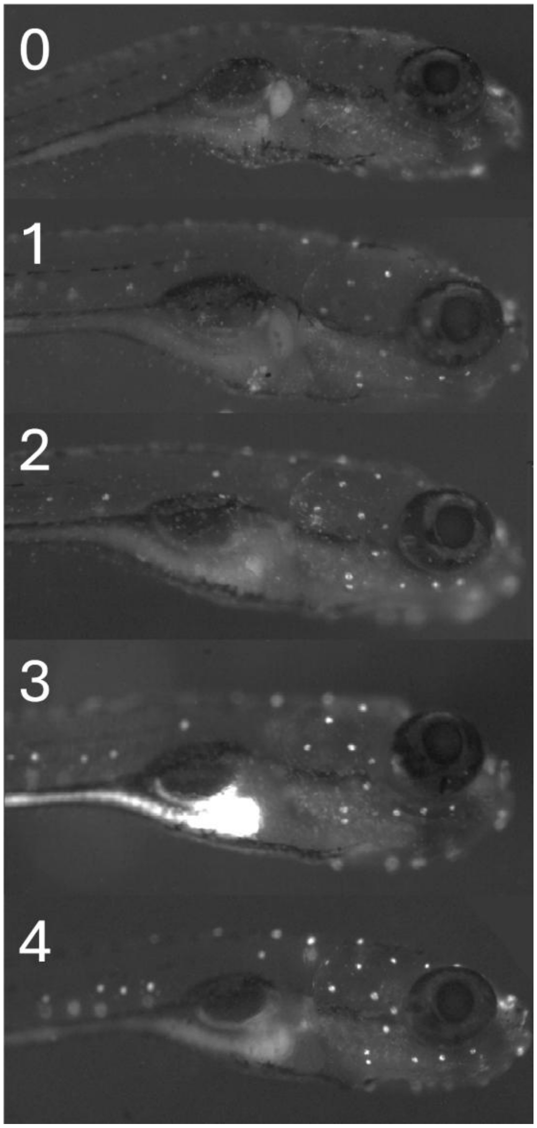


Figure S1. Representative images illustrating the neuromast scoring scale (0–4), depicting progressive levels of hair cell damage and fluorescence intensity from complete loss (0) to intact neuromasts (4).

**Compound Toxicity in Zebrafish**

To establish safe working concentrations for otoprotection assays, dose–response studies were conducted to evaluate compound-associated toxicity in larval zebrafish. Both acute (1-hour), and chronic (24-hour) exposures were assessed to determine effects on neuromast integrity and overall organismal viability. The zebrafish lateral line system provides a sensitive in vivo platform for detecting both hair cell–specific and systemic toxicity.

In acute exposure assays, compound 28510 exhibited a clear dose-dependent reduction in neuromast quality, with significant toxicity observed between 50 and 100 µM. Similarly, compound 663865 showed dose-dependent toxicity, with significant decreases in neuromast integrity occurring between 100 and 150 µM. In contrast, compound 27671 did not exhibit detectable toxicity, with neuromast quality remaining unaffected at concentrations up to 250 µM (Figure S2A).

In chronic exposure assays, 28510 exhibited reduced toxicity relative to continuous exposure, with significant decreases in neuromast quality observed only at higher concentrations (50–100 µM). Compound 663865 produced modest toxicity at ≥150 µM, while 27671 remained well tolerated with no significant effects on neuromast integrity (Figure S2B).

Overall, these findings demonstrate that 28510 and 663865 exhibit dose- and exposure-dependent toxicity profiles in zebrafish, whereas 27671 displays a markedly improved safety profile. These data informed the selection of dosing ranges for subsequent otoprotection assays and highlight the importance of balancing efficacy with tolerability in lead compound optimization.

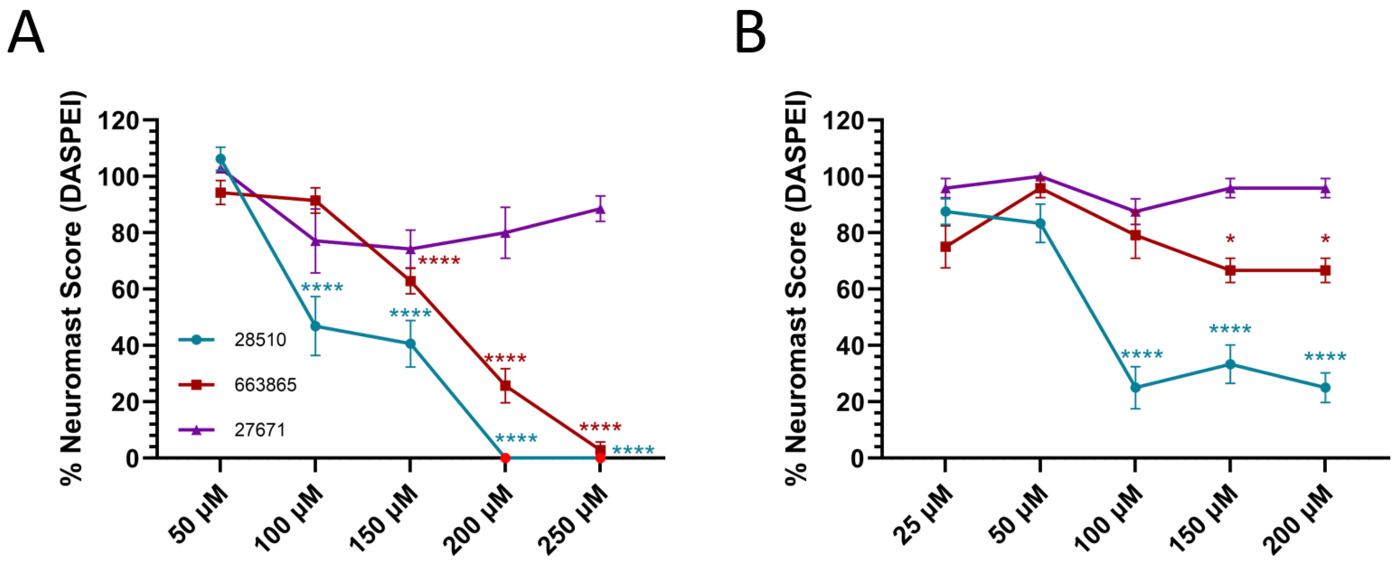


Figure S2. Dose–response curves of structural analogs showing effects on zebrafish neuromast quality following A) acute treatment, B) chronic treatment. Values represent mean whole-fish neuromast quality expressed as a percentage of untreated controls ± SEM (*n* = 9-12). Statistical significance was determined by one-way ANOVA with Tukey's post hoc test (*p<0.05, **p<0.01, ***p<0.001, **p<0.0001), with asterisks indicating comparisons versus untreated controls.

**Drug Combinations**

To evaluate potential synergistic or additive effects, combinations of lead compounds were tested in zebrafish hair cell protection assays. Specifically, combinations of 28510+663865 and 663865+27671 were assessed to determine whether reduced individual doses could preserve otoprotective efficacy while minimizing dose-dependent toxicity. Combination approaches are commonly employed in otoprotection studies to enhance efficacy while reducing adverse effects associated with single-agent treatments (Owens et al., 2009; Coffin et al., 2010).

**Combination of 28510 + 663865**

The combination improved the tolerated dose range. While 28510 alone reduced neuromast quality at higher concentrations (e.g., 100 μM), the combined treatment (50+50 μM; total 100 μM) did not adversely affect neuromast integrity. Toxic effects emerged only at higher combined doses (75+75 μM; total 150 μM) (Figure S3), suggesting modest improvement in tolerability under chronic treatments.

Functionally, the 28510+663865 combination produced significant otoprotection in acute gentamicin and neomycin assays; however, these effects were not superior to those observed with either compound alone (Figure S4A). In chronic AG exposure assays, the combination restored neuromast integrity to untreated control levels in kanamycin and streptomycin conditions at 10+10 μM (total 20 μM), but these effects were not significantly different from those achieved by 28510 alone (Figure S4B). Overall, these results indicate limited evidence of synergy, with modest improvements in tolerability but no enhancement of otoprotective efficacy beyond that of the lead compound.

**Combination of 27671 + 663865**

In contrast, the combination of 27671 and 663865 exhibited a distinct interaction profile. In dose–response assays, the combination produced greater toxicity at 50+50 μM (total 100 μM) than either compound alone at equivalent doses (Figure S3), further indicating reduced tolerability.

Despite these limitations, the 27671+663865 combination demonstrated enhanced otoprotective effects under certain conditions. In acute assays, the combination provided significant protection against gentamicin and neomycin at 25+25 μM (total 50 μM), with modest improvement over individual compounds, particularly in neomycin exposure (Figure S5A).

In chronic AG exposure assays, the combination improved neuromast quality in gentamicin- and streptomycin-treated groups, although not to levels comparable to untreated controls. Notably, at 5+5 μM (total 10 μM), the combination provided significant protection against gentamicin, exceeding the effects of either compound alone (Figure S5B), suggesting a potential low-dose synergistic interaction under specific conditions.

Overall, while combination strategies revealed context-dependent interactions, neither pairing consistently demonstrated robust synergy across assays. The 28510+663865 combination modestly improved tolerability without enhancing efficacy, whereas the 27671+663865 combination exhibited both increased toxicity and limited, condition-specific gains in otoprotection.

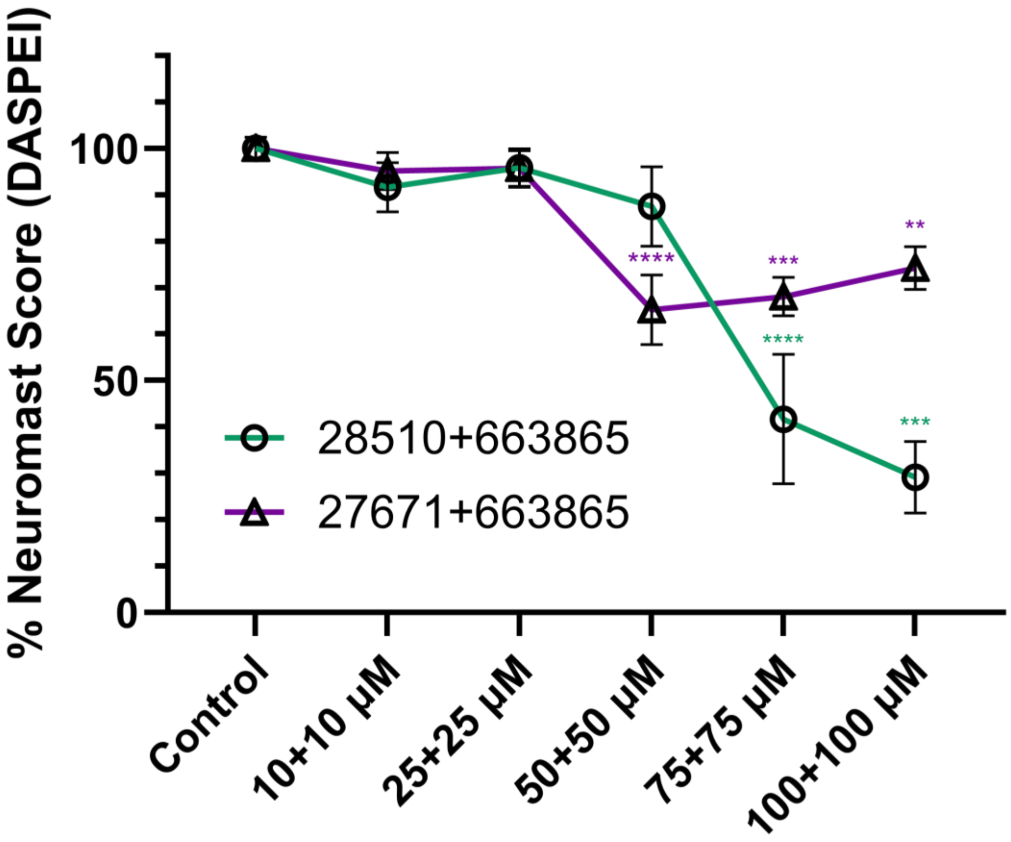


Figure S3. Dose–response curve showing the effects of structural analog combinations on zebrafish neuromast integrity. Values represent mean whole-fish neuromast quality expressed as a percentage of untreated controls ± SEM (*n* = 9-12). Statistical significance was determined by one-way ANOVA with Tukey's post hoc test (*p<0.05, **p<0.01, ***p<0.001, **p<0.0001) versus untreated controls.

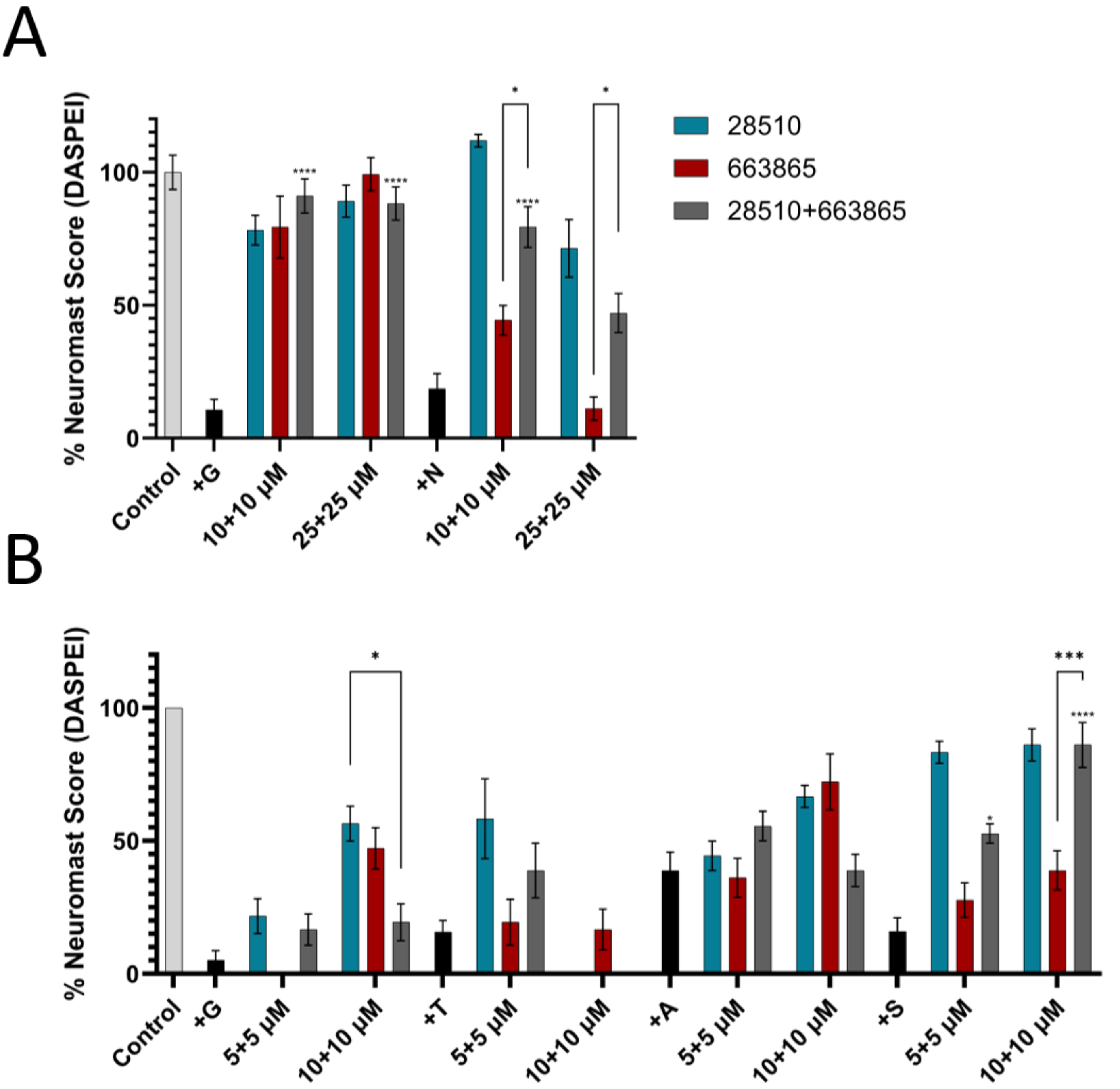


Figure S4. Comparison of otoprotective effects of 28510, 663865, and their combination (28510+663865). (A) Acute protection against 110 μM gentamicin and 50 μM neomycin. (B) Chronic protection against 200 μM gentamicin, 5 mM kanamycin, 1 mM tobramycin, 500 μM streptomycin, and 10 mM amikacin. Combination doses are presented as paired concentrations (e.g., 5+5 μM) and correspond to equivalent total doses of individual compounds (e.g., 10 μM), with 10+10 μM corresponding to 25 μM and 25+25 μM to 50 μM. Values represent mean whole-fish neuromast quality expressed as a percentage of untreated controls ± SEM (*n* = 9-12). Statistical significance for otoprotection was determined by one-way ANOVA with Tukey's post hoc test versus AG-treated controls. Comparisons between treatment groups were analyzed by two-way ANOVA with Tukey's post hoc test (*p<0.05, **p<0.01, ***p<0.001, **p<0.0001).

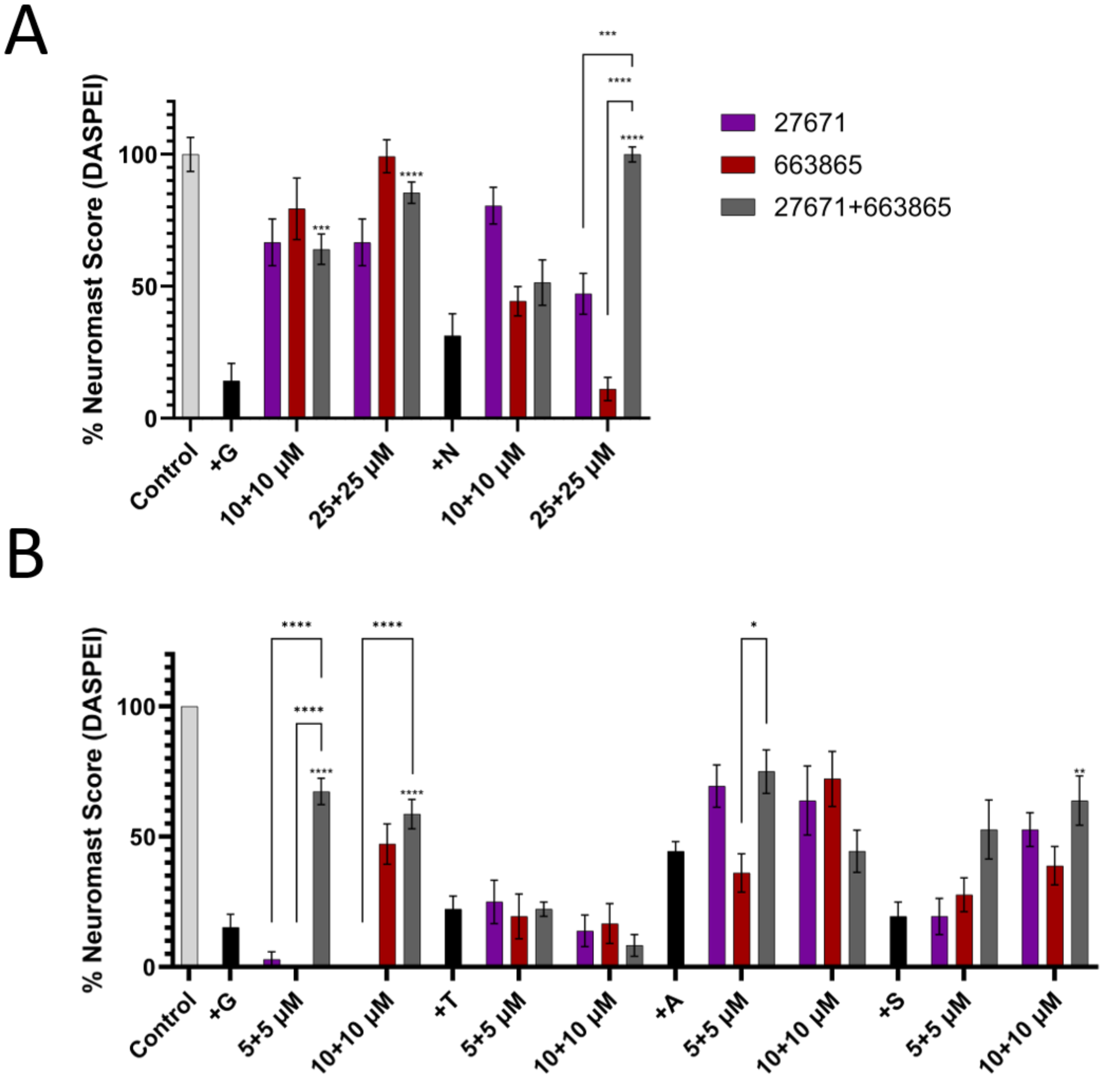


Figure S5. Comparison of otoprotective effects of 27671, 663865, and their combination (27671+663865). (A) Acute protection against 110 μM gentamicin and 50 μM neomycin. (B) Chronic protection against 200 μM gentamicin, 5 mM kanamycin, 1 mM tobramycin, 500 μM streptomycin, and 10 mM amikacin. Combination doses are presented as paired concentrations (e.g., 5+5 μM) and correspond to equivalent total doses of individual compounds (e.g., 10 μM), with 10+10 μM corresponding to 25 μM and 25+25 μM to 50 μM. Values represent mean whole-fish neuromast quality expressed as a percentage of untreated controls ± SEM (*n* = 9-12). Statistical significance for otoprotection was determined by one-way ANOVA with Tukey's post hoc test versus AG-treated controls. Comparisons between treatment groups were analyzed using two-way ANOVA with Tukey’s post hoc test (*p< 0.05, **p<0.01, ***p<0.001, **p<0.0001).
